# Rapid metabolic and behavioural maladaptations following short-term obesogenic diet withdrawal during post-weaning development in male Wistar rats

**DOI:** 10.64898/2026.03.12.710091

**Authors:** Breno Picin Casagrande, Vitória Rios Beserra, Alessandra Mussi Ribeiro, Luciana Pellegrini Pisani, Debora Estadella

## Abstract

**Background:** Obesogenic diets (ODs) are known to trigger metabolic and inflammatory disturbances. However, the effects of short-term OD withdrawal on systemic and neuroinflammatory parameters remain unclear.

**Objectives:** This study investigated the short-term effects of OD withdrawal on metabolic, inflammatory, and anxiety-like behaviours in young male Wistar rats.

**Methods:** Three-week-old male Wistar rats were fed either a control (Ct, n=5) or high-sugar/high-fat (HSHF) diet for 14 days. Animals in the HSHF group were further divided into no-withdrawal (NWt, n=5) and withdrawal (Wt, n=5) groups, where Wt received a control diet for 48 hours. Food intake, body mass, adiposity, serum metabolic parameters, hepatic energy stores, inflammatory markers (serum, liver, hypothalamus, hippocampus, mesenteric fat), and oxidative stress markers in the hippocampus were measured. Elevated plus maze and open field were used to characterize behavioural phenotype.

**Results:** OD intake significantly increased caloric intake, visceral adiposity, hepatic glycogen, and TAG levels. The 48-hour withdrawal reduced TAG, induced hyperinsulinemia and hypoglycaemia, and heightened inflammation in mesenteric fat, serum, and the hippocampus. Oxidative stress markers (SOD and MDA) increased in the hippocampus, correlating with elevated serum corticosterone and heightened anxiety-like behaviour in the Wt group compared to the other groups.

**Conclusion:** Short-term withdrawal after only two weeks of OD intake exacerbates systemic and neuroinflammation, hippocampal oxidative stress, and anxiety-like behaviours, indicating rapid negative responses to dietary transition. These findings identify short-term dietary withdrawal during early development as a distinct and biologically active phase, rather than an immediate recovery period.

## Introduction

ODs are characterized by high levels of sugar, fat, or a combination of both, are well-known for their negative effects on several respects of general health [1,2]. Long term improvements in diet can provide significant health benefits, but short-term metabolic and behavioural consequences of shifting to a healthier diet show mixed results [1,3,4]. Despite the expected general recovery at any point after dietary improvement, it does not immediately nor necessarily happen, especially at the short-term [5,6].

The abrupt removal of energy-dense, palatable and obesogenic diets can elicit metabolic and behavioural responses that resemble addictive substance withdrawal, including reduced food intake, craving, heightened stress responses, and increased systemic inflammation [5–8]. These responses are thought to arise from metabolic and neuroendocrine shifts, but the precise mechanisms remain unclear. These outcomes are particularly concerning during early life, a critical period for brain development and susceptibility to dietary stressors [9–11].

Understanding the underlying neurobehavioral mechanisms of OD withdrawal is essential, particularly given its association with anxiety-like behaviours. Although the nutritional components – sugar, fat and palatability – do not have drug-like effects *per se*, the feeding behaviour and its ambience lead to pro-reward and anti-stress effects that are self-reinforcing in a drug-like manner. Similar patterns are seen in other behaviours, where physiological and behavioural adaptations sustain the habit and contribute to relapse [12–15]. In the context of ODs, these processes may also shape the response to withdrawal, particularly in the short term.

ODs are known to disrupt metabolic control and affect behaviour [1,2,16], it increases adiposity and promotes low-grade inflammation and oxidative stress [17–19]. These changes contribute to insulin resistance and lipid accumulation, in the liver, excess carbohydrate intake can increase de novo lipogenesis and fat deposition [20,21]. These effects are partly driven by oxidative stress caused by reactive oxygen species (ROS), which, while essential for cellular signalling in moderate amounts, become damaging when unbalanced [22]. Excess ROS leads to lipid peroxidation, protein carbonylation, and irreversible cellular damage, increasing the risk of chronic diseases [23]. Adipose tissue expansion further increases the risk of insulin resistance, due to the release of free fatty acids, ROS, and inflammatory cytokines [24,25].

The brain is markedly vulnerable to these disturbances due to its high metabolic demand and limited antioxidant capacity [26]. Oxidative stress and neuroinflammation activate microglia and astrocytes, leading to sustained cytokine production and disruption of neuronal function [27,28]. Within the brain, the hippocampus is especially sensitive to these insults, while it plays a central role in memory and emotional regulation [29]. Cytokines such as TNFα and IL6 impair synaptic plasticity and neurogenesis, partly through disruption of BDNF signalling [30–32]. Consistent with this, increased hippocampal inflammation and oxidative damage have been linked to anxiety-like behaviour in rodent models [33–35].

The post-weaning period is marked by rapid brain development, including synaptic remodelling and maturation of neuroendocrine systems [36,37]. During this stage, the brain is more sensitive to environmental and metabolic challenges [38,39], thus, nutritional stressors in early development can produce more pronounced and persistent consequences [40]. While OD withdrawal has been studied in adult animals [8,41], much less is known about its effects during this developmental window, which are likely more affected by short-term negative responses and could impact adherence to healthier diets [42].

In this study, we examined the short-term effects of OD withdrawal in post-weaning male Wistar rats. Animals were exposed to an OD for only two weeks, followed by a brief 48-hour withdrawal period. We hypothesized that even brief OD exposure, followed by withdrawal, would disrupt metabolic homeostasis and increase anxiety-like behaviour. This design allows the characterization of a short dietary transition during post-weaning development, in which abrupt withdrawal after a brief obesogenic exposure may be associated with concurrent metabolic, inflammatory, oxidative, and behavioural alterations.

## Methods

### Ethics and sample size calculations

The ethics committee of the Federal University of Sao Paulo (n° 9512270320) approved all procedures. National and international ethical principles and guidelines for animal research were followed. We calculated the sample size with G*Power software (version 3.1.9.7) considering a high effect size (f=1 or r²=0.50), with power (1−β) at 80%, and alpha at 5% as the convention. The calculation returned a total of 15 animals. The effect size used for the calculations was extracted from previous data and considering the primary outcome “anxiety-like behaviour” on the elevated plus maze [8] and hippocampal inflammation [43].

### Study design

Post-weaning male Wistar rats (3 weeks old) were used in this study. The animals underwent adaptation to the vivarium for 7 days. At 4 weeks of age (28d), the rats were assigned to the treatment groups and received the control or obesogenic diet and water for 14 days *ad libitum* (Table 1). The obesogenic diet, a high-sugar/high-fat diet (HSHF), was adapted from our previous HSHF diet model [44] to meet the AIN-93 protein recommendations for the growth period [45]. The HSHF diet was prepared by adding sweetened condensed milk (Nestlé) and lard (Sadia, São Paulo, Brazil) to ground control chow. To meet nutritional recommendations [45], the formulation was adjusted with casein (Labsynth, Diadema, Brazil), soybean oil (Bunge, Gaspar, Brazil), and a vitamin and mineral mix (RHOSTER, Araçoiaba da Serra, Brazil). Additionally, to reflect the excess sodium intake observed globally, approximately double the recommended levels [2,46], sodium chloride was added to the diet (11.6 g/kg, corresponding to 4.5 g sodium) to achieve a twofold increase in sodium content in the final formulation. This amount was calculated based on the sodium content of the control diet (2700 mg/kg). While excess fat and sugar intake are the most studied components of ODs, high sodium intake is also a major contributor to diet-related disease, largely driven by processed foods [46].

**Table 1.** Diet composition.

| **Components (g/100g)** | **High-sugar/high-fat diet** | **Control diet** |
| --- | --- | --- |
| Control diet | 285.48 | 1000.00 |
| Sweetened condensed milk | 386.57 | 0.00 |
| Lard | 79.06 | 0.00 |
| Casein | 79.06 | 0.00 |
| Sucrose | 114.19 | 0.00 |
| Soybean oil | 10.76 | 0.00 |
| Mineral mix (AIN-93) | 25.43 | 0.00 |
| Vitamin mix (AIN-93) | 8.14 | 0.00 |
| Butylhydroxytoluene | 0.11 | 0.00 |
| Sodium chloride | 11.62 | 0.00 |
| **Sodium** | **5.27** | **2.70** |
| **Nutrients (g/1000g)** |  |  |
| Carbohydrates | 493.32 | 583.30 |
| Sucrose | 326.80 | 0 |
| Fiber | 72.43 | 253.70 |
| Protein | 138.93 | 209.70 |
| Lipids | 133.42 | 44.40 |
| Added saturated fat | 50.16 |  |
| **Energy (%)** |  |  |
| Carbohydrates | 48.94 | 51.56 |
| Sucrose | 38.00 | 0 |
| Protein | 16.15 | 32.81 |
| Lipids | 34.91 | 15.63 |
| Saturated fat | 13.12 | 0 |
| **Energy (Kcal/1000g)** | **3440.08** | **2556.80** |

Group allocation was performed using stratified randomization based on absolute body mass and its variation during the adaptation period. The animals were assigned to either the control diet group (n=5) or the high-sugar, high-fat (HSHF) diet group (n=10). The control group (Ct) was fed the control diet for 14 days, while the HSHF group was divided into a no-withdrawal subgroup (NWt, n=5), which was fed the HSHF diet for 14 days, and a withdrawal subgroup (Wt, n=5), which was fed the HSHF diet for 14 days followed by the control diet for 2 days. As a result, the withdrawal group remained under experimental conditions for an additional 48 hours compared to the other groups, which represents a potential time-related confound. The animals were group housed (5 animals/cage).

Food intake was measured every 2–3 days by calculating the difference between the amount of food provided and the sum of the remaining food and any food scattered in the bedding. It was expressed as grams consumed per animal per 24h per 100g of body mass. Energy intake was calculated according to food intake and diet composition (Table 1) and is reported as kcal consumed per animal per 24h per 100g of body mass. Body mass was evaluated at the beginning, each week, and at the end of the protocol. Also, we measured it every day of the second week and during the 2 days of withdrawal. The percentage of body mass gained was calculated at each point accordingly.

At the end of the protocol behavioural tests were conducted. After a fasting period of 6–8 hours, the animals were anaesthetized with isoflurane (1.25ml/L) in a sealed transparent plastic box and euthanized by decapitation after confirming complete anaesthesia. Trunk blood, hypothalamus, hippocampus, liver, and visceral adipose tissues – mesenteric, epididymal, and retroperitoneal – were collected. The tissue masses were verified, and they were reported relative to body mass. In all analysis, the investigator was blinded to the treatment conditions.

### Behavioural testing

The animals were fasted for 1 hour before testing while they adapted to the room. The tests began at 8h00, they were conducted in the same conditions across the whole experiment. The room had dimmed lights and controlled noise. The apparatus was cleaned with 70% and 5% alcohol and was let dry after each run.

The open filed (OF) and the elevated plus maze (EPM) were used to determine anxiety-like behaviour. The OF consisted of a 60 cm circular arena subdivided in central, 30 cm diameter, surrounded by a peripheral zone, with 15 cm. Each animal was placed individually in the centre of the apparatus and recorded for 5 minutes. Total lines crossed, time in the centre, and lines crossed in the peripheral zone were evaluated with ANY-maze (v7.4, Stoelting Co, USA). The EPM was set 50 cm above the floor. The apparatus consisted of four intercalated arms – two open and two enclosed –, each measuring 60 cm × 15 cm. The centre of the apparatus was a square of complementary dimensions (15 cm × 15 cm). Each one of the animals was placed in the centre facing the same open arm and it was recorded for 5 min with free access to the entire maze. Entries, distance, time, average speed, and immobility time in each section of the maze were evaluated with ANY-maze (v7.4, Stoelting Co, USA).

To characterize the anxiety-like behaviour, the time in the centre of the OF, and the parameters of the EPM, the percentage of entries, time, and distance on the open arms [47] in relation to the total number of entries, time, and distance on the whole apparatus, open arms, enclosed arms, and centre (*e.g.*, entries on the open arms/sum of entries in the open arms, enclosed arms, and centre). Additionally, we evaluated risk assessment via protected and unprotected head dips, which was done manually by two independent blinded researchers. Then the Principal Component Analysis (PCA) approach was used for further data interpretation being able to unify measures from both tests in a single component.

### Serum parameters

Trunk blood collected during the euthanasia after 6-8 hours fasting was separated into serum (10 min centrifugation at 5,000 ×*g* at 4°C). Serum glucose (#133, Labtest, Brazil), insulin (80-INSMR-CH01, STELLUX®, ALPCO, US), triacylglycerol (TAG) (#87, Labtest, Brazil), total cholesterol (#76, Labtest, Brazil), and HDL-cholesterol (#13, Labtest, Brazil) were determined with colourimetric kits following the manufactures instructions. HOMA-IR index and n-HDL/HDL cholesterol ratio were calculated.

### Hepatic glycogen and lipids

Hepatic glycogen levels were determined by adapting the micro-method proposed by Balmain, Biggers, and Claringbold (1956) as previously standardized by our group [48,49]. Hepatic lipids were extracted following the Folch et al (1957) method [50]. From those lipids, we quantified TAG, cholesterol, and HDL-cholesterol using the above-reported colourimetric kits (Labtest, Brazil).

### Inflammation

Inflammatory cytokines were assessed in the hypothalamus, liver, hippocampus, serum, and mesenteric fat. Epididymal and retroperitoneal fat deposits were not evaluated since we found no important cytokine alterations in these tissues in previous work [44]. The serum was prepared for the analysis as previously described [43]. Tissues were homogenized in ice-cold lysis buffer (100 mM Tris-HCl, 10 mM sodium pyrophosphate, 10 mM sodium orthovanadate, 2 mM phenylmethylsulfonyl fluoride, 0.04% bovine lung aprotinin; 10 mM EDTA, 100 mM sodium fluoride, 1% Triton) and centrifuged at 20,000 ×*g* for 40min at 4°C. The supernatant was transferred to another microtube, and protein content was measured with Bradford reagent [51]. The concentrations of IL6 (#DY506, R&D Systems, US), IL10 (#DY522, R&D Systems, US), IL1β (#DY501, R&D Systems, US), and TNFα (#DY510, R&D Systems, US) were evaluated in the supernatant following the manufacturer’s instructions.

### Oxidative Status

Hippocampi samples were homogenized in 0.2M PBS pH 7.4 and centrifuged at 4°C for 15 min at 5,000×g. The supernatant was used to estimate the activity of catalase (CAT), superoxide dismutase (SOD), and malondialdehyde (MDA) concentrations, the pellet was used to estimate the concentration of carbonylated proteins (CBP).

CAT activity was estimated as described by Góth (1991), which is based on hydrogen peroxide neutralization, the reaction was carried out for 3 minutes and interrupted by the addition of ammonium molybdate. The calculations were carried out to estimate the activity of catalase as hydrogen peroxide units consumed per minute, and it was adjusted by the protein content of the sample [52]

SOD content was estimated by the autoxidation of pyrogallol and continued superoxide-dependent reduction of 3-(4,5-dimethyl-thiazol-2-yl) 2,5-diphenyl tetrazolium bromide (MTT) to formazan. This reaction was incubated for 15 minutes at 37°C and was interrupted by the addition of dimethyl sulfoxide (DMSO). The results are shown as SOD units per mg of protein [53]

MDA concentration was estimated by the thiobarbituric acid reactive substance (TBARS) assay. Samples were incubated in a water bath at 90° for 40 minutes, we performed an alcoholic extraction with n-butanol, and its phase was recovered by centrifugation for 10 minutes at 5000×g. The results are shown as µM/mg protein [54,55].

Protein carbonylation was estimated by derivatization with 2,4-dinitrophenylhydrazine (DNPH) in a 2M HCl solution in the dark. The proteins were precipitated with a 10% trichloroacetic acid (TCA) solution and washed twice with equal parts ethyl acetate and ethanol solution. Between the washes, the samples were centrifuged (10 minutes, 4°C, 10,000×g), and the supernatant was removed. In the final step, a 6% SDS solution was added, the sample was homogenized and centrifuged once again (10 minutes, 23°C, 10,000×g), and the supernatant was used for the readings. A blank was run without the DNPH addition alongside each sample. After the calculations, the result is expressed in nmol/mL [56]

## Statistical analysis

Data were represented as mean ± standard deviation, while are presented as median with interquartile range, with individual data points overlaid in graphical representations. For all analysis, jamovi software was used (version 2.6.2) alongside GAMLj3 module (3.4.2) [57]. Due to the small sample size, no outlier exclusion procedures were applied, outliers were identified and *winsorization* was employed to preserve robustness and stabilize variance [58]. We used Generalized linear models (GzLM) and Generalizes Estimated Equations (GEE) to analyse single and multiple observation data, respectively. Distribution was selected based on data type and AIC. Holm’s stepwise correction was used in all cases. To verify if the food and calorie intake and was different upon withdrawal in the Wt group, the Z-test was performed. Pearson’s correlations were used to determine relationship between variables. Fisher r-to-z transformation was also employed to determine differences between observed correlations. The Principal Component Analysis (PCA) was used to combine EPM variables to determine anxiety-like behaviour. Statistical significance was set at 5% as convention and the effect sizes are reported. For GZLM and GEE, χ² values, p-value, residual degrees of freedom (rDf), and r² were reported; for between-group comparisons, p-value and Hedges’ *g* were reported. The sample size calculations were performed considering a power level (1 – β error) of 80%. To attend the matter of the sample size and concerns about the validity of the findings, it is expected that the r² is greater than 0.5 for GLZMs (single observation) and r² greater than 0.066 for GEEs (repeated observations), considering the required Cohen’s f (1 and 0.266).

## Results

### Food intake, body mass, and adiposity

Food and calorie intake are displayed in Figure 1.A-B. Because animals were housed in a single cage per group, we could not analyse the group-by-time interaction, however, it is possible to determine the effects of group and time separately.

**Figure 1.**
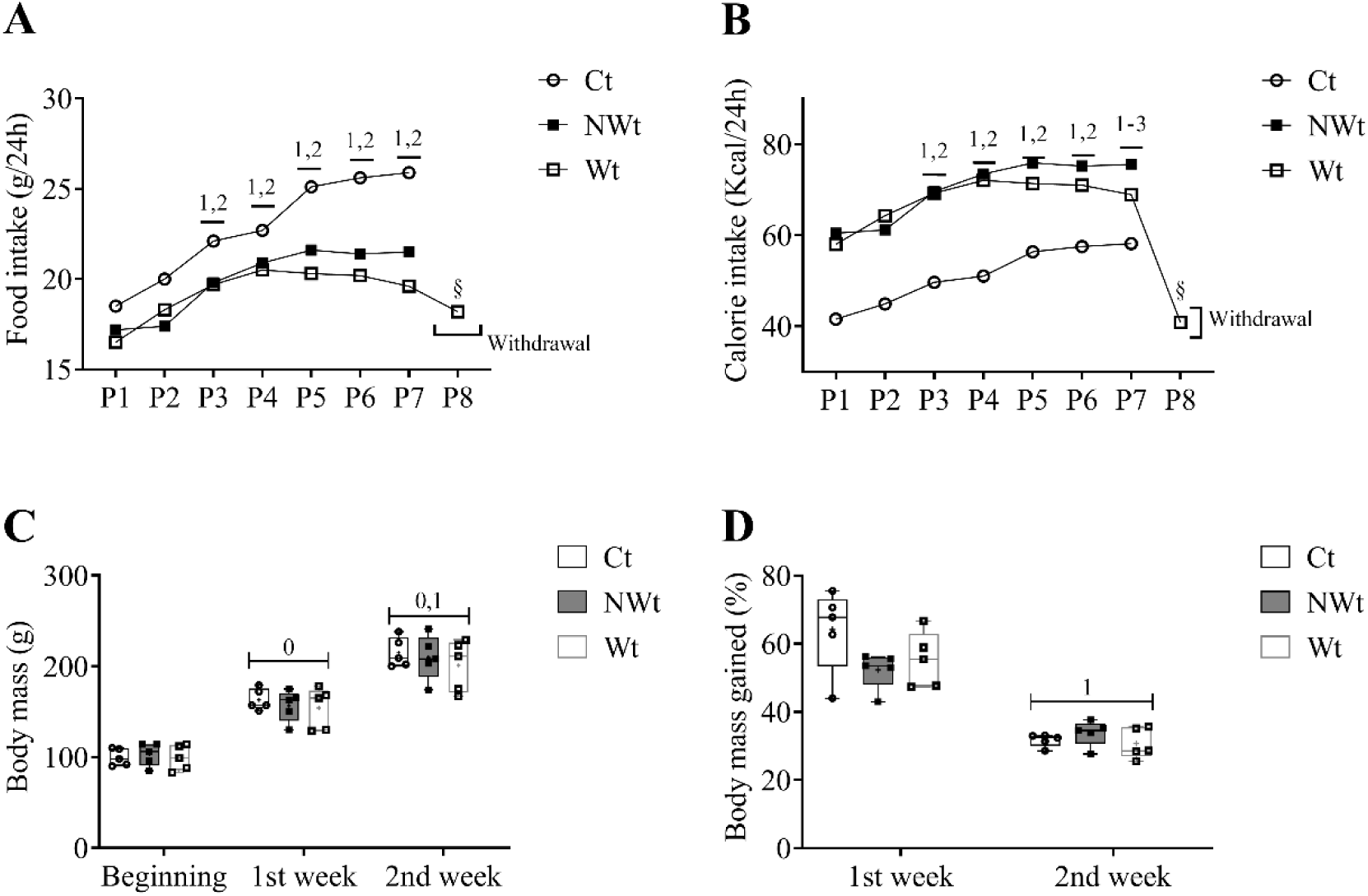
Food and calorie intake, body mass, and percentage body mass gained over time. (A) Food intake consumed (g) per 24h per animal, the intake was measured every 2-3 days (n=1 cage/group); P8 is withdrawal period. (B) Calorie intake consumed (kcal) per 24h per animal, the intake was measured every 2-3 days (n=1 cage/group); P8 is withdrawal period. (C) Body mass (g) per group each week (n=5/group). (D) Body mass gained (%) per group each week (n=5/group). (A, B) numbers 1-3 indicate p<0.05 comparing to respective period (P) (1 to P1, 2 to P2) § indicates p<0.05 for one-tailed Z-test compared to Wt intake. (C, D). numbers 0, 1 indicate p<0.05 compared to the beginning (0) and 1^st^ week (1).

Food (χ²=9.67, p < 0.001, rDf=6, r²=0.92) and calorie (χ²=6.93, p = 0.002, rDf =6, r²=0.39) intake increased with time. Post-hoc comparisons are indicated in Figure 1.A-B. The HSHF-fed groups consumed less chow than the Ct group Ct vs. NWt p < 0.001, g = 2.57, Ct vs. Wt p < 0.001, g = 3.43). Conversely, they consumed more calories than Ct (Ct vs. NWt p < 0.001, g = −7.55, Ct vs. Wt p < 0.001, g = −6.52). At period 8, the withdrawal for Wt, the food intake dropped in both kcal (two-tail Z-test p < 0.001) and grams (two-tail Z-test p = 0.042).

Initial and final body masses were similar among groups (Table 2 and Supplementary Table 1). Although, body mass increased in all groups in the first and second week compared to the previous (χ²=506.90, p < 0.001, rDf =24, r²=0.84; Beginning vs. 1^st^ p < 0.001, g = −3.33; Beginning vs. 2^nd^ p < 0.001, g = −6.39; 1^st^ vs. 2^nd^ p < 0.001, g = 2.98) (Figure 1.C). However, the percentage of gain decreased from the first to the second week (χ²=129.76, p < 0.001, rDf = 12, r²=0.84; 1^st^ vs. 2^nd^ p < 0.001, g = 2.99). There were no differences across groups in either body mass (χ²=0.39, p = 0.682, rDf = 12) or body mass gain (χ²=1.42, p = 0.278, rDf = 12) progressions (Figure 1.D).

**Table 2.**
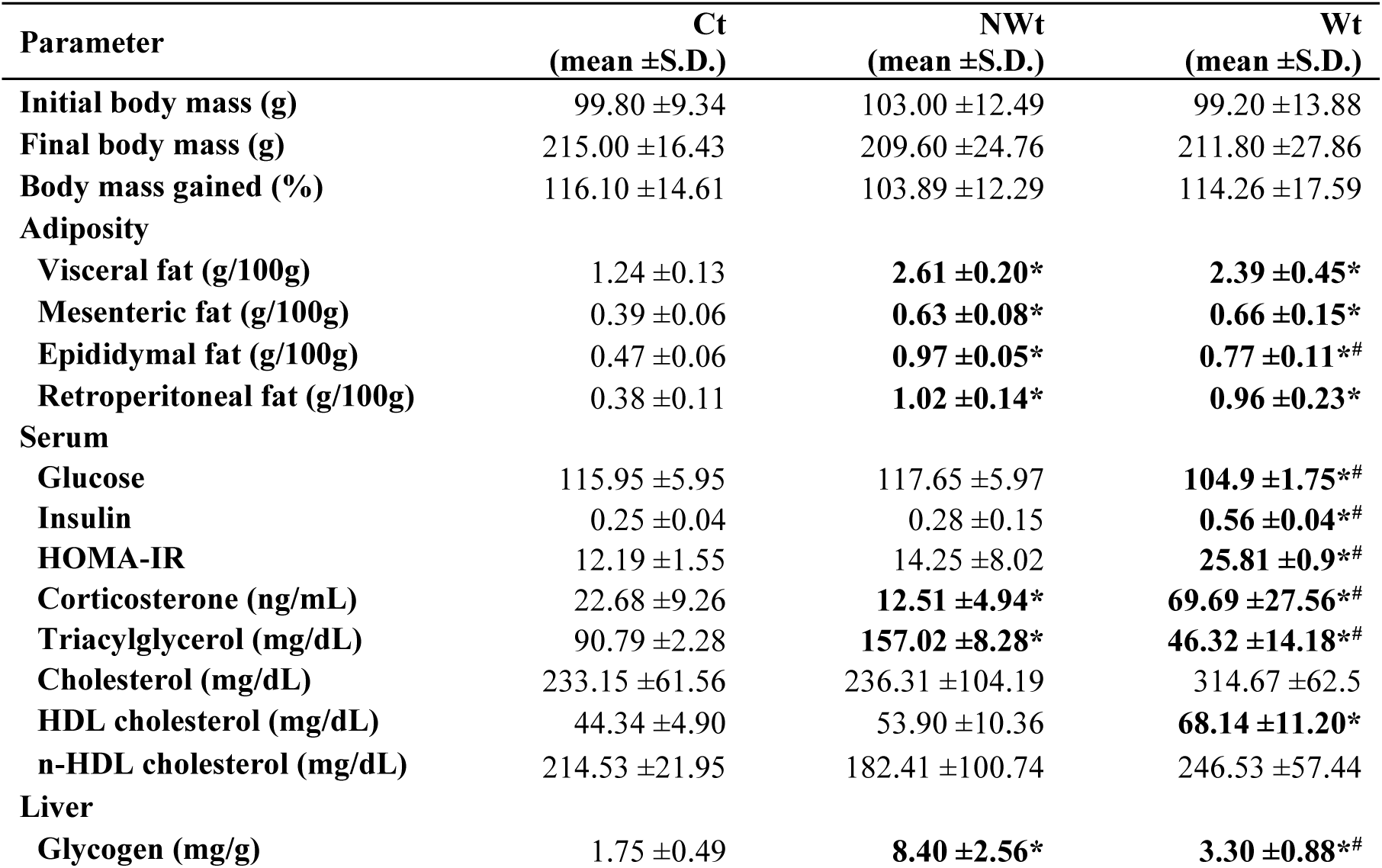
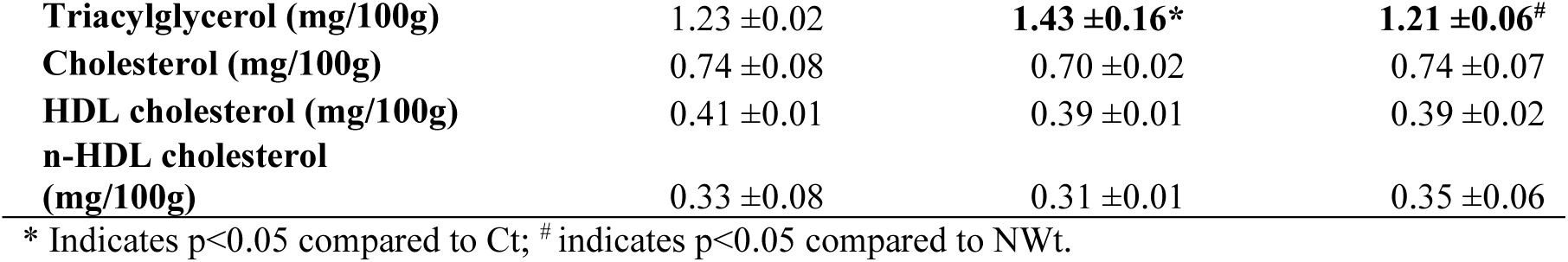
Body mass, adiposity, serum parameters, and metabolic parameters.

Overall body mass gained was also similar among groups, yet visceral adiposity (g/100g) was different (Table 2). It was higher in HSHF-fed groups compared to Ct (Ct vs. NWt p < 0.001, g = −3.60; Ct vs. Wt p < 0.001, g = −3.24). Mesenteric, epididymal, and retroperitoneal fat deposits were increased in NWt and Wt groups compared to Ct (Mesenteric Ct vs. NWt p = 0.008, g = −2.72; Ct vs. Wt p = 0.004, g = −1.92; Epididymal Ct vs. NWt p < 0.001, g = −8.86; Ct vs. Wt p < 0.001, g = −2.65; Retroperitoneal Ct vs. NWt p < 0.001, g = −4.18; Ct vs. Wt p < 0.001, g = −2.58) (Table 2 and Supplementary table 1). Wt group presented intermediate values for epididymal fat, lower than NWt (p = 0.002, g = 1.86).

### Serum and hepatic parameters

Serum glucose was lower in the Wt group compared to Ct and NWt (Ct vs. Wt p = 0.001, g = 2.10; NWt vs. Wt p < 0.001, g = 2.40). Alongside, Wt had higher insulin (Ct vs. Wt p = 0.002, g = −2.08; NWt vs. Wt p = 0.006, g = −2.31) and HOMA-IR index (Ct vs. Wt p = 0.002, g = −8.71; NWt vs. Wt p = 0.004, g = −1.64). Table 2, Supplementary Table 1)

NWt group had higher serum TAG compared to the other groups (Ct vs. NWt p < 0.001, g = −8.82 NWt vs. Wt p < 0.001, g = 7.71). Wt presented lower TAG than the Ct group (Ct vs. Wt p < 0.001, g = 3.54). HDL cholesterol was higher in Wt compared to Ct (Ct vs. Wt p = 0.005, g = −2.22). No alterations were seen in total nor n-HDL cholesterol. (Table 2, Supplementary Table 1)

Hepatic glycogen was higher in the NWt group compared to the other groups (Ct vs. NWt p < 0.001, g = −2.91; NWt vs. Wt p < 0.001, g = 2.15). The Wt also presented higher glycogen than the Ct group (Ct vs. Wt p = 0.003, g = −1.76). Hepatic TAG was also higher in NWt compared to the other groups (Ct vs. NWt p = 0.015, g = −1.42; NWt vs. Wt p = 0.010, g = 1.52). (Table 2, Supplementary table 1)

### Inflammation

In the serum, IL6, and IL10 were not different among groups. There was an increase in serum TNFα in the Wt group compared to Ct (Ct vs. Wt p = 0.010, g = −1.73). Serum IL1β was higher in the NWt group compared to the other groups (Ct vs. NWt p = 0.003, g = −2.04; NWt vs. Wt p = 0.008, g = 1.76). (Figure 2.A, Supplementary Table 2)

**Figure 2.**
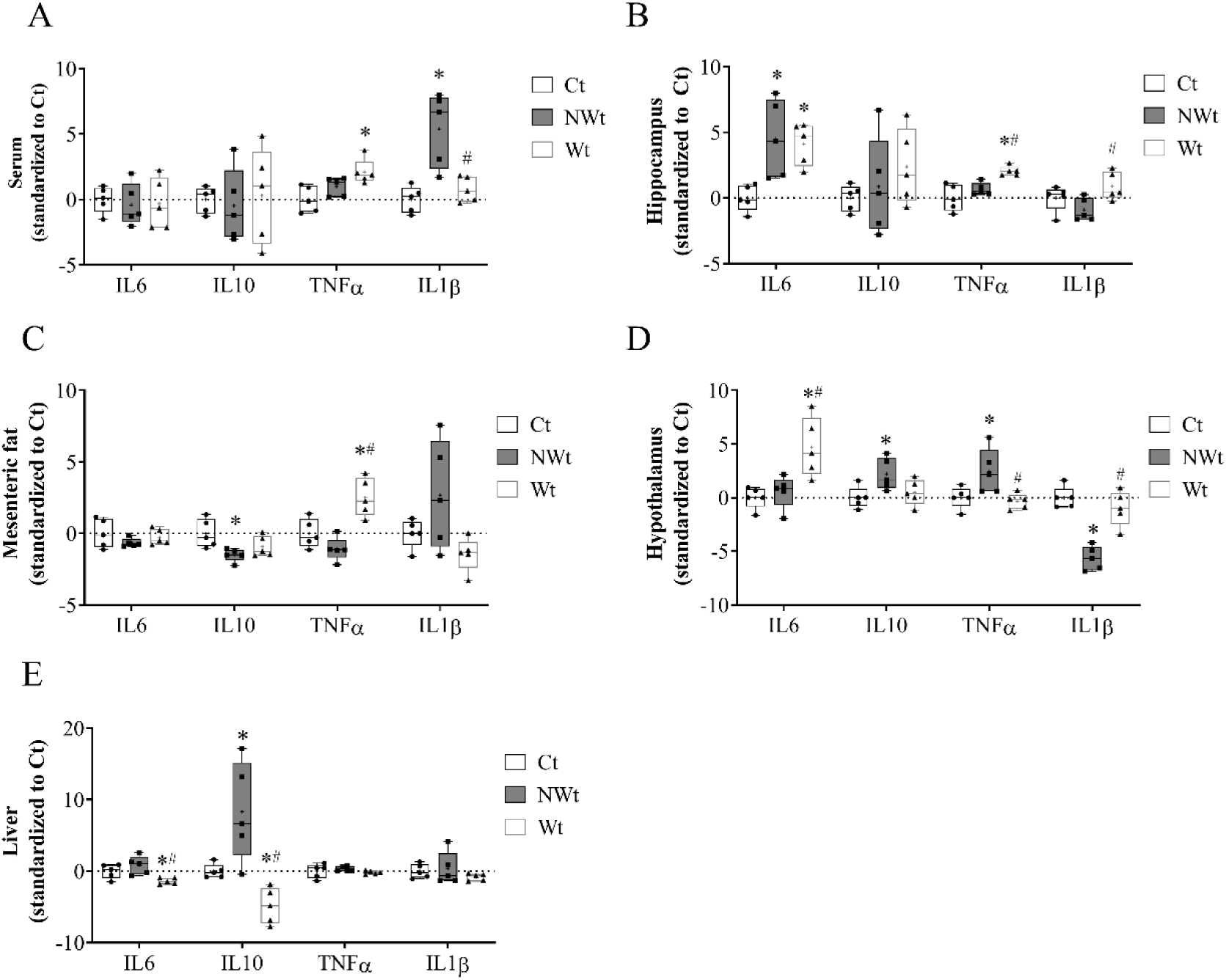
Graphical representation standardized values of cytokines. (A) Serum; (B) Hippocampus; (C) Mesenteric fat; (D) Hypothalamus; (E) Liver. * Indicates p<0.05 compared to Ct; ^#^ indicates p<0.05 compared to NWt. Cytokine values are shown as Z-scores to facilitate comparison across tissues and cytokines with different concentration ranges; absolute values are reported in Supplementary Table 3.

In the hippocampus, NWt and Wt presented higher IL6 levels compared to Ct (Ct vs. NWt p = 0.007, g = −1.65; Ct vs. Wt p = 0.007, g = −2.49). TNFα was higher in the Wt group compared to the other groups (Ct vs. Wt p = 0.004, g = −2.18; NWt vs. Wt p = 0.037, g = −2.55). Wt showed an increase in IL1β compared to NWt (NWt vs. Wt p = 0.040, g = -1.49). (Figure 2.B, Supplementary table 2)

In mesenteric fat, IL10 was lower in NWt compared to Ct (p = 0.023, g = 1.56). TNFα was higher in the Wt group compared to the other groups (Ct vs. Wt p = 0.007, g = −1.72; NWt vs. Wt p < 0.001, g = −2.61) (Table 3). (Figure 2.C, Supplementary table 2)

**Table 3.** Hippocampal oxidative stress.

| Parameter | Ct<br>(mean ±S.D.) | NWt<br>(mean ±S.D.) | Wt<br>(mean ±S.D.) |
| --- | --- | --- | --- |
| SOD (units/mg of protein) | 209.11 ±12.99 | 217.69 ±31.69 | **334.50 ±52.86\*<sup>#</sup>** |
| CAT (H<sub>2</sub>O<sub>2</sub> units consumed /min/mg of protein) | 0.34 ±0.13 | 0.35 ±0.09 | 0.35 ±0.03 |
| MDA (μmol/mg of protein) | 0.17 ±0.02 | 0.17 ±0.01 | **0.23 ±0.05\*<sup>#</sup>** |
| CBP (mmol/mL) | 1.60 ±0.43 | **0.65 ±0.21\*** | **0.51 ±0.14\*** |

SOD: superoxide dismutases; CAT: catalase; MDA: malondialdehyde; CPB: carbonylated protein. \*
Indicates p<0.05 compared to Ct; <sup>#</sup> indicates p<0.05 compared to NWt.

In the hypothalamus, IL6 was higher in Wt compared to Ct and NWt (Ct vs. Wt p = 0.006, g = −1.48; NWt vs. Wt p = 0.010, g = −1.81). IL10 was higher in NWt compared to Ct (Ct vs. NWt p = 0.049, g = −1.42). TNFα was also increased in NWt group (Ct vs. NWt p = 0.032, g = −1.21; NWt vs. Wt p = 0.022, g = 1.44). Conversely, IL1β presented an opposite response, NWt showed lower concentration than Ct and Wt (Ct vs. NWt p < 0.001, g = −4.42; NWt vs. Wt p < 0.001, g = −2.69). (Figure 2.D, Supplementary Table 2)

In the liver, Wt presented a decrease in IL6 and IL10 content compared to Ct (IL6 p = 0.031, g = 1.48; IL10, p = 0.047, g = 2.07) and NWt group (IL6, p = 0.005, g = 1.94; IL10, p < 0.001, g = 2.04). NWt showed higher concentrations than Ct group (p = 0.016, g = −1.36). (Figure 2.E, Supplementary table 2)

### Hippocampal oxidative status

Hippocampal SOD was higher in the Wt group compared to Ct and NWt (Ct vs. Wt p < 0.001, g = −2.65; NWt vs. Wt p < 0.001, g = −2.24) and the same pattern was seen for MDA (Ct vs. Wt p = 0.022, g = −1.23; NWt vs. Wt p = 0.013, g = −1.50). CBP was lower in NWt and Wt groups compared to Ct (Ct vs. NWt p = 0.001, g = 2.29; Ct vs. Wt p < 0.001, g = 2.82). (Table 3, Supplementary Table 2)

### Behavioural testing

Total lines crossed on the OF was higher in the NWt (p = 0.006, g = −2.93) group but decreased upon withdrawal (p = 0.006, g = 2.86). The same effect was observed with lines crossed on the peripheral zone (Ct vs. NWt p= 0.012, g= −2.53; NWt vs. Wt p = 0.002, g = 2.99). Time in the centre of the OF was lower on the Wt group compared to NWt (p = 0.027, g = −1.36) (Figure 3.A, Supplementary Table 4). No differences were seen in zone crossing and entries in both regions (data not shown).

**Figure 3.**
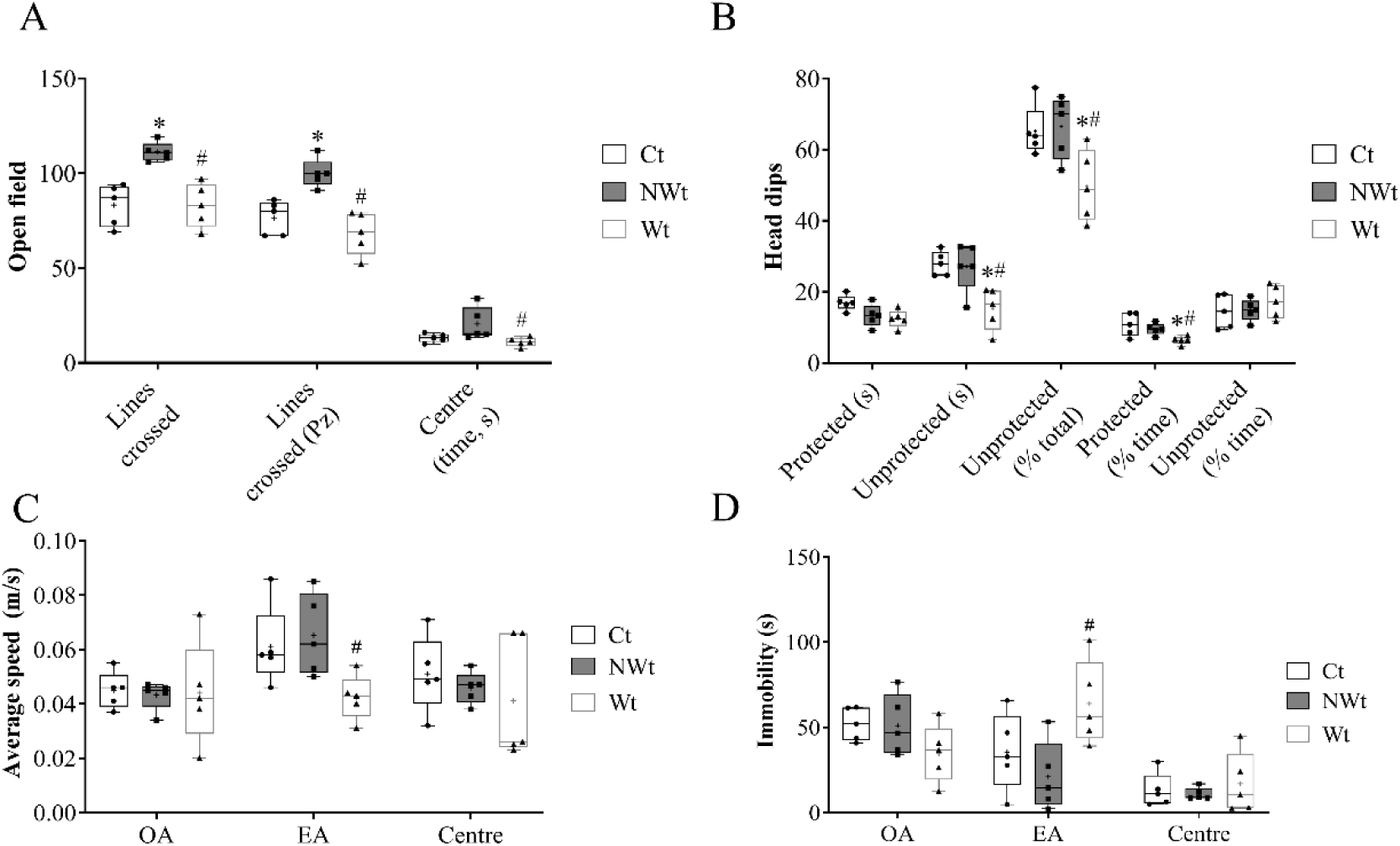
Graphical representation of behavioural parameters. (A) open field lines crossed and time in the centre; (B) risk assessment in the elevated plus maze as protected and unprotected head dips; (C) average speed in the sections of the elevated plus maze; (D) immobility time in the sections of the elevated plus maze. * Indicates p<0.05 compared to Ct; ^#^ indicates p<0.05 compared to NWt. (OA) open arms; (EA) enclosed arms.

Total distance in the EPM was lower in the Wt group compared to both Ct and NWt (Average (± SD), Ct 2.10 ±0.17; NWt 2.06 ±0.17; Wt 1.22 ±0.25) (χ² = 62.09, p < 0.001, rDf = 2, r² = 0.84; Ct vs. Wt p < 0.001, g = 3.56; NWt vs. Wt p < 0.001, g = 3.53). In the open arms (OA) of the EPM, the Wt group presented lower percentages of entries (Ct vs. Wt p = 0.001, g = 2.04; NWt vs. Wt p = 0.003, g = 2.41), time (Ct vs. Wt p = 0.011, g = 1.41; NWt vs. Wt p = 0.007, g = 2.80), and distance (Ct vs. Wt p = 0.001, g = 2.68; NWt vs. Wt p = 0.002, g = 1.64) than the Ct and NWt groups. (Figure 4.A, Supplementary Table 4). No differences were seen in the enclosed arms (EA). (Figure 4.B, Supplementary Table 4). To determine whether reduced exploration reflected anxiety-like behaviour or locomotor impairment, we analysed average speed and immobility across EPM. No differences were observed in either parameter in the OA or centre. In the EA, the Wt group exhibited lower average speed and increased immobility; however, these effects were associated with low r² and insufficient statistical power (Figure 3.C-D, Supplementary Table 4).

**Figure 4.**
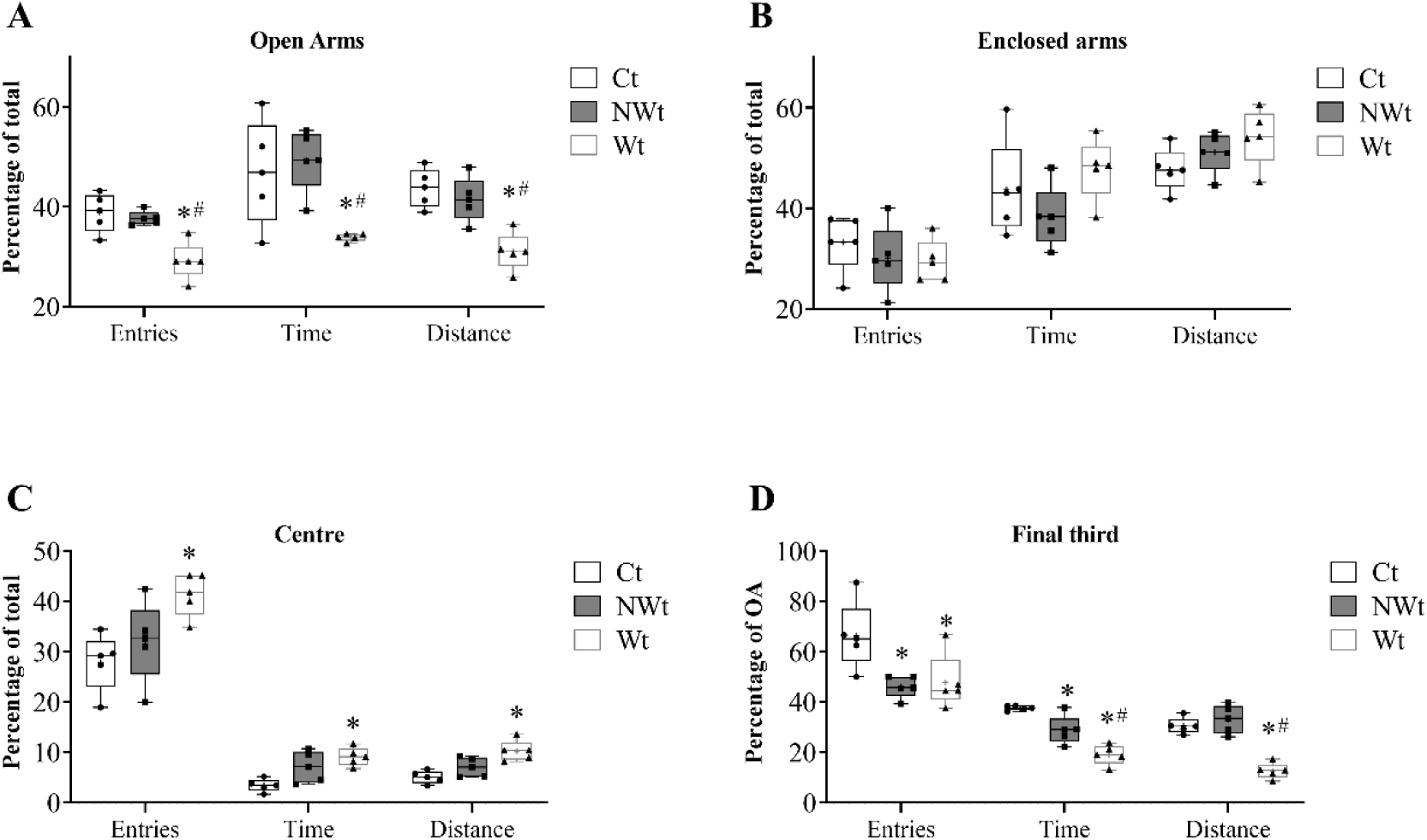
Graphical representation of elevated plus maze. (A) entries (% total), time (% total), and distance (% total) on the open arms; (B) entries (% total), time (% total), and distance (% total) on the enclosed arms; (C) entries (% total), time (% total), and distance (% total) on the centre; (D) entries (% open arms), time (% open arms), and distance (% open arms) on the final third of the open arms. * Indicates p<0.05 compared to Ct; ^#^ indicates p<0.05 compared to NWt.

In the centre (Ce), the NWt group showed higher percentages of time (p = 0.023, g = −1.26) than Ct group. While the Wt group presented higher percentages of entries (p = 0.029, g = −2.16), time (p = 0.009, g = −2.92), and distance (p = 0.004, g = −2.48) compared to the Ct group. (Figure 4.C, Supplementary Table 4)

In the final third of the OA, the NWt and Wt groups showed lower percentage of entries (Ct vs. NWt p = 0.032, g = 1.63; Ct vs. Wt p = 0.038, g = −1.21) and time than Ct group (Ct vs. NWt p = 0.030, g = −1.72; Ct vs. Wt p < 0.001, g = −5.23). Whereas Wt groups showed a further decrease compared to NWt in the percentage of time (p = 0.004 g = 1.64). Wt groups also had lower percentage of distance travelled in the final third compared to both Ct and NWt (Ct vs. Wt p < 0.001, g = 4.59; NWt vs. Wt p < 0.001, g = 3.64). (Figure 4.D, Supplementary Table 3)

The percentage of time in the OA was negatively correlated with the percentage of time spent in the EA (r= −0.786, r²=0.61, p < 0.001), while the percentage of distance on the OA was negatively correlated with the percentage of distance on the EA (r= −0.792, r²=0.627, p < 0.001). The correlations were adjusted for group conditions.

Risk assessment was also different among groups. Wt group showed lower head dips in the open arms, as unprotected head dips, compared to both Ct (p = 0.011, g = 2.38) and NWt (p = 0.012, g = 1.66). While there was a trend (p = 0.053, g = 1.78) towards lower protected head dips compared to Ct group. Wt group also showed a lower percentage of unprotected head dips, as a percentage of total head dips, compared to both Ct (p = 0.033, g = 1.60) and NWt (p = 0.033, g = 1.66). As the time in each section of the EPM was different among groups, we calculated the percentage of time on head dips in the protected and unprotected sections. With this approach, Wt group presented lower protected head dips considering time under protected conditions – at the centre or enclosed arms – compared to both Ct (p = 0.012, g = 2.07) and NWt (p = 0.033, g = 1.65), but no differences were seen for unprotected head dips in the same conditions. (Figure 3.B, Supplementary Table 4)

For the anxiety-like behaviour determined via PCA, named as ANX, its component comprised the following variables and had the following component loadings: entries, distance, and time in the OA (entries: −0.79; distance: −0.94; time: −0.87), distance and time in the EA (distance: 0.77; time: 0.81) related to totals on the apparatus; protected (−0.57) and unprotected head dips (−0.94); and time in the centre of the OF (−0.45). The greater the score, the greater the anxiety-like behaviour. ANX (Average (± SD), Ct −0.59 ±0.74; NWt −0.49 ±0.62; Wt 1.08 ±0.62) was higher in the Wt group compared to both Ct (p = 0.005, g = −2.21) and NWt (p = 0.006, g = −2.29).

### Correlations for Hypothalamic Cytokines and Energy homeostasis

Several correlations were seen among hypothalamic cytokines and energy homeostasis parameters evaluated. For all groups, hypothalamic IL6 was correlated with insulin (r = 0.77, p < 0.001, 95% CI 0.43 to 0.92), HOMA-IR (r = 0.74, p = 0.002, 95% CI 0.36 to 0.91), corticosterone (r = 0.65, p = 0.009, 95% CI 0.20-0.87), serum glucose (r = − 0.53, p = 0.043, 95% CI −0.02 to −0.82), and serum TAG (r = − 0.54, p = 0.037, 95% CI −0.04 to −0.82). As OD intake altered these parameters, we evaluated if the observed correlations differed when analysing them separately. For the HSHF-fed groups, the correlations were mostly maintained apart from with corticosterone and glycaemia (insulin r = 0.74, p = 0.014, 95% CI 0.21 to 0.94; HOMA-IR r = 0.69, p = 0.027, 95% CI 0.11 to 0.92; corticosterone r = 0.61, p = 0.062; serum glucose r = − 0.54, p = 0.107; serum TAG r = − 0.69, p = 0.028, 95% CI −0.92 to −0.10).

The HSHF-fed groups also presented opposing correlations of these parameters with hypothalamic TNFα (insulin r = − 0.82, p = 0.004, 95% CI −0.96 to −0.39; HOMA-IR r = − 0.79, 95% CI −0.95 to −0.39, p = 0.006; corticosterone r = − 0.72, p = 0.018, 95% CI −0.93 to −0.17; serum glucose r = 0.64, p = 0.044, 95% CI 0.02 to 0.91; serum TAG r = 0.69, p = 0.027, 95% CI 0.11 to 0.92). To confirm the significance of the opposing nature of the relationships, we applied the Fisher r-to-z transformation. All correlations were significantly different (insulin z = 3.94, p = 0.0001; HOMA-IR z = 3.59, p = 0.0003; corticosterone z = 3.02, p = 0.0025; serum glucose z = −2.55, p = 0.0108; serum TAG z = −3.17, p = 0.0015).

While when all groups are taken into account, only hypothalamic TNFα correlation with corticosterone and TAG are seen (insulin r = − 0.50, p = 0.054; HOMA-IR r = − 0.49, p = 0.061; corticosterone r = − 0.57, p = 0.027, 95% CI −0.84 to −0.08; serum glucose r = 0.43, p = 0.107; serum TAG r = 0.65, p = 0.008, 95% CI 0.21 to 0.87).

### Correlations for Anxiety-like behaviour

The relevant correlations found are presented on table 4. It is worth highlighting that parameters indicating a more anxious-like phenotype were correlated with an inflammation hallmark, TNFα, in the serum, hippocampus, and mesenteric fat. Additionally, they were correlated with the oxidative status of the hippocampus, being directly associated with the increase in MDA and SOD. Likewise, it was directly correlated with serum corticosterone, a key marker of stress system activation.

**Table 4.** Correlations between inflammation, oxidative stress, and the open arms parameters.

|  | Serum<br>Cort. | Serum<br>TNFα | Hepatic<br>IL6 | Ht<br>IL6 | Hc<br>TNF | M.FAT<br>TNFα | Hc<br>SOD | Hc<br>MDA |
| --- | --- | --- | --- | --- | --- | --- | --- | --- |
| **Centre (OF)** |  |  |  |  |  |  |  |  |
| Time (s) |  |  | **0.696\*** |  |  | **- 0.514\*** |  |  |
| **Open Arms (EPM)** |  |  |  |  |  |  |  |  |
| Entries (% total entries) | - 0.801* | - 0.623* |  | - 0.699* | - 0.515* | - 0.765* | - 0.727* | - 0.526* |
| Time (% total time) | - 0.685* |  | 0.723* | - 0.704* |  | - 0.589* | - 0.753* |  |
| Distance (% total distance) | - 0.675* |  | 0.662* | - 0.815* |  | - 0.577* | - 0.636* |  |
| **Final third of the OA (EPM)** |  |  |  |  |  |  |  |  |
| Time (% open arms) | - 0.656* | - 0.694* |  | - 0.710* | - 0.718* | - 0.567* | - 0.718* | - 0.561* |
| Distance (% open arms) | - 0.762* |  | 0.812* | - 0.794* | - 0.591* | - 0.691* | - 0.834* | - 0.661* |
| **Unprotected head dips (EPM)** | **- 0.735\*** | **- 0.558\*** | **0.616\*** | **- 0.852\*** |  | **- 0.667\*** | **- 0.630\*** |  |
| **ANX (PCA)** | **0.704\*** |  | **0.709\*** | **0.839\*** |  | **0.645\*** | **0.743\*** |  |

OF: open field; Cort: corticosterone; Hc: hippocampal; Ht: hypothalamic; M.FAT: mesenteric fat; SOD: superoxide dismutase; MDA: malondialdehyde; ANX: anxiety-like behaviour; PCA: Principal Component Analysis. \* indicates p<0.05

## Discussion

In this study, we evaluated the metabolic and behavioural effects of withdrawing from an obesogenic diet after two weeks of intake in a post-weaning model. In the present post-weaning model, withdrawal was associated with rapid metabolic disruption and the emergence of anxiety-like behaviour within 48h. These results indicate that metabolic and behavioural systems in early life can respond strongly to abrupt dietary change, even after brief exposure. Importantly, these effects did not occur in isolation. These outcomes occurred concurrently, suggesting that multiple metabolic, inflammatory, oxidative, and behavioural responses were engaged in parallel during abrupt dietary transition, rather than implying a single causal pathway. The temporal convergence of these outcomes suggests a coordinated physiological response to abrupt dietary transition rather than independent alterations across systems.

### Metabolic and Inflammatory Effects of Two-Week Obesogenic Diet

Food intake (g) was higher in the Ct group, but calorie intake was higher in groups consuming the HSHF diet. Despite higher caloric intake over two weeks, body mass did not differ between groups. This likely reflects the rapid growth phase at this age [59], and suggests changes in body composition, supported by higher visceral adiposity (Table 2).

Two weeks of HSHF intake caused dyslipidemia, increased hepatic glycogen, hepatic TAG deposition, and increased hypothalamic and serum inflammation. These findings partially contrast with previous studies in which one- and four-week OD feeding did not consistently increase hepatic TAG [60]. In addition, while four weeks of OD intake elevated inflammation in the liver and mesenteric adipose tissue, the two-week protocol was insufficient to induce similar changes [44]. Adolescent rats show greater hepatic lipogenic capacity, including sustained increases in fatty acid synthase and stearoyl-CoA desaturase activity after short-term fructose feeding [61], but they may also show reduced baseline and stimulated pro-inflammatory responses [62]. Younger animals may therefore be more sensitive to early metabolic alterations, including hepatic glycogen accumulation and TAG deposition [49].

The magnitude of changes after two weeks of OD intake indicates rapid metabolic disruption in early life, including a 110% increase in visceral adiposity, a 16% increase in hepatic lipid accumulation, and a 380% increase in hepatic glycogen. Although hepatic inflammation was not detected, the accumulation of visceral fat and hepatic glycogen likely reflects early adaptive responses to caloric excess. This supports the view that short-term OD exposure can produce meaningful metabolic changes, even if inflammatory changes are not yet present. While some reports suggest inflammation precedes and may contribute to hepatic TAG accumulation [44,63], our pattern is consistent with the natural history of fatty liver [64].

The absence of a direct anxiogenic effect after 14 days of OD intake is consistent with prior work. Vega-Torres and colleagues reported that 11 days of a high-fat diet during adolescence did not affect anxiety-like behaviour in the EPM [65], although findings across studies can be heterogeneous [5,66], and are likely driven by differences in diet composition, such as fat and sugar content.

### Acute Metabolic and Behavioural Shifts upon Diet Withdrawal

Upon withdrawal, animals voluntary decreased calorie intake, which may reflect lower preference for non-palatable foods and/or the lower calorie density of the control diet [5,67]. Visceral adiposity remained elevated compared to Ct, indicating that short-term withdrawal did not reverse the adiposity increase induced by the HSHF diet.

After withdrawal, animals showed metabolic and behavioural changes consistent with a shift from an anabolic to a catabolic state. Compared with prior findings in older male Wistar rats exposed to four weeks of HSHF feeding [43,44,68], the two-day withdrawal after two-week OD intake showed a distinct metabolic response. This likely reflects differences in developmental stage, duration of exposure, and available energy stores, rather than a directly greater effect. [49,59,69,70].

A briefer OD intake provides less opportunity for energy storage, and thus weaker “protection” to low energy availability states, which may be the cause for an amplified negative physiological impact to the OD removal. Consistently, the 48h withdrawal did not restore metabolic homeostasis and was associated with poor glucose and lipid handling, with a state of hypoglycaemia, increased insulin and HOMA-IR, reduced circulating TAG, and persistent visceral adiposity.

As expected, anxiety-like behaviour increased after OD withdrawal. In the open field, withdrawal animals differed from NWt, and in the EPM they showed reduced open arm exploration and altered head dip measures compared with Ct and NWt. Effect sizes were large and comparable to prior work. The overall component (ANX) shows a consistent response. In the present study, the model effect size was r²=0.63 (Bootstrap 95% CI 0.265 to 0.836), while a previous study reported r²=0.47. Between-group comparisons also showed Hedges’ g values within the reported range. Compared to control groups, we observed g=−2.21 (95% CI −3.38 to −0.54), while a previous study reported g=−1.50; compared to animals still on OD, we observed g=−2.29 (95% CI −3.91 to −0.59), while a previous study reported −2.19 [8].

In adult mice, the trajectory appears different. Garcia-Serrano et al. reported that long-term obesogenic diet exposure induces metabolic and behavioural impairments that are largely reversed after prolonged diet normalization [71]. This suggests that in mature systems, diet-induced alterations can be reversible with sufficient recovery time. Our findings are not directly comparable due to differences in developmental stage and recovery duration, but together they indicate that response dynamics depend on age and timescale. The stronger response observed after short-term exposure may be related to lower energy storage (e.g., visceral fat) compared to longer feeding protocols, which could increase sensitivity to abrupt reductions in caloric intake.

Reduced open arm exploration can be confounded by locomotor changes, but this is unlikely here. Average speed and immobility time in the open arms were unchanged, suggesting preserved motor capacity in the anxiogenic context. Modest changes in enclosed arms (reduced speed and increased immobility) had insufficient statistical power and may reflect increased passivity (e.g., coping or risk avoidance) rather than generalized hypoactivity. Neophobia is another potential confound, but prior work shows increased anxiety-like behaviour after sucrose withdrawal in paradigms that minimize neophobia [72]. Similarly, Sharma and colleagues reported increased anxiety-like behaviour one day after high-fat diet withdrawal, but not after removal of a low-fat diet [73]. Accordingly, the rodents were previously familiar with the control diet, which they were fed during acclimatization.

### Neuroinflammation, Oxidative Stress, and Emergence of Anxiety-Like Behaviour

NWt animals showed higher serum IL1β, a systemic contributor to atherosclerosis [74], but lower hypothalamic IL1β than Ct. In contrast, hypothalamic TNFα was elevated. Different hypothalamic inflammatory patterns are associated with distinct energy homeostasis profiles. Stimuli from saturated fatty acids and nutrient excess are linked to diet-induced obesity, whereas stimuli derived from plasma cytokines/LPS are associated with sickness behaviour and hypophagia [75]. TNFα can modulate insulin and leptin signalling and exert anorexigenic effects [76], while IL1β regulates cephalic insulin release[77].

Following withdrawal, Wt animals showed a spike in serum TNFα compared to Ct and increased hypothalamic IL6 compared to both groups, consistent with prior findings [68]. Hypothalamic IL6 has been shown to increase fatty acid oxidation in skeletal muscle [78], which may reflect a response to reduced energy availability. IL6 can also suppress feeding and improve peripheral glucose homeostasis [79], consistent with the observed reduction in intake and metabolic changes.

Hypothalamic IL6 correlated positively with serum insulin, insulin sensitivity, and corticosterone, and negatively with glycaemia and serum TAG. These associations are consistent with IL6 being linked to coordinated central and peripheral responses during withdrawal, but they do not establish causality. The positive correlations may reflect an adaptive response to reduced energy availability, while the negative correlations may reflect compensatory changes in peripheral energy utilization, potentially involving IL6-related increases in fatty acid oxidation [78].

Within the HSHF-fed groups, most correlations were sustained, and additional associations emerged: hypothalamic TNFα correlated negatively with insulin (r = − 0.82, p = 0.004), HOMA-IR (r = − 0.80, p = 0.006), corticosterone (r = − 0.72, p = 0.018), alongside positive correlations with glycaemia (r = 0.64, p = 0.045) and serum TAG (r = 0.69, p = 0.027). These patterns highlight the complexity of cytokine interactions in the hypothalamus during withdrawal. While TNFα is often linked to inflammation and insulin resistance [80], the direction of associations here may reflect context-dependent regulation during acute dietary transition. IL6 and TNFα patterns suggest potentially opposing roles in response to OD withdrawal. These correlations were observed primarily during the withdrawal condition and were not uniformly present during continued HSHF feeding.

The hippocampus was selected due to its established role in stress regulation and anxiety-related behaviour [29,30,81], and its vulnerability to oxidative stress and inflammation [26]. After OD withdrawal, hippocampal pro-inflammatory cytokines and oxidative stress markers were elevated compared to Ct and NWt. TNFα can disrupt glutamatergic signalling and reduce synaptic plasticity [82,83], and elevated IL6 can exacerbate neuroimmune activation [84]. These mechanisms could contribute to altered hippocampal function relevant to anxiety-like behaviour, but this remains a hypothesis based on known biology rather than direct evidence in this study. The findings are consistent with literature linking neuroinflammation to reduced BDNF expression and impaired neurogenesis and synaptic connectivity [32,85], processes implicated in anxiety disorders.

OD withdrawal also induced hippocampal oxidative stress, with elevated MDA and SOD compared to Ct and NWt. MDA reflects lipid peroxidation and oxidative damage to neuronal membranes, while increased SOD likely reflects a compensatory response to increased ROS. Hippocampal oxidative stress and inflammatory markers, as well as serum corticosterone correlated with anxiety-like behaviours in the EPM (Table 4), supporting an association between these biological changes and the behavioural phenotype. Increased serum corticosterone is consistent with stress-axis activation, which can worsen hippocampal oxidative stress and impair neuroplasticity. These results align with mechanisms implicated in anxiety-related and neuropsychiatric conditions [86], but further work is needed to test causal pathways.

### Implications for Dietary Transition and Intervention Strategies

Negative consequences of OD can manifest within 24 hours [87] and persist as long as tested, although outcomes vary with sex, age, and diet composition and can change over time. Hence, short-term withdrawal should not be interpreted as the beginning of the recovery phase. Instead, the present findings indicate that early withdrawal represents a distinct transitional state marked by physiological stress, which differs from both continued OD exposure and long-term dietary normalization.

The 48-hour withdrawal protocol produced both expected and unexpected outcomes. While prior studies report metabolic and behavioural impacts of OD exposure, we did not expect such pronounced negative consequences after such short exposure. These findings suggest an early onset of mechanisms underlying the anxiogenic response to OD withdrawal, which indicates that prolonged exposure may not be necessary for addiction-related behaviours to emerge in this early-life stage.

At the same time, long-term dietary improvements provide substantial benefits, highlighting the need for supporting strategies during early phases of dietary change. Pharmacotherapy and bariatric surgery can be effective in adults but are not primarily recommended for younger populations [88,89]. Therefore, developing and evaluating non-pharmacological and behavioural interventions tailored to early life and to the early withdrawal period are needed.

## Limitations

This study should be interpreted as characterizing short-term withdrawal responses during early development rather than long-term outcomes or permanent alterations. First, the model targets the early post-weaning period, when neural and metabolic systems are still maturing. The responses likely reflect interactions between acute metabolic effects and developmental plasticity and may not generalize to fully mature organisms. Second, the withdrawal period was limited to 48 hours to capture early-phase responses; it does not address longer-term recovery trajectories and is not directly comparable to studies using extended normalization periods. Third, animals were housed in a single cage per group, introducing potential cage-level effects, including social hierarchy and shared microenvironment, which cannot be excluded post hoc. However, the consistency of outcomes across metabolic, inflammatory, and behavioural endpoints suggests the phenotype is unlikely to be driven by a single confounder. Fourth, despite being based on sample size calculations, the small group sizes (n=5 per condition) limit generalization. Fifth, only male animals were included, limiting extrapolation across sexes given known sex-specific differences in responses to obesogenic diets. Finally, the dietary model was adapted to approximate key features of ultra-processed diets (high fat, sugar, and sodium). While this increases translational relevance, it reduces standardization relative to purified diets and may limit direct comparison with defined formulations.

## Statements and Declarations

### Funding

This work was supported by ‘São Paulo Research Foundation’ (FAPESP #2021/02325-0) and the ‘Coordination for the Improvement of Higher Education Personnel (CAPES Brazil - Financial Code 001). LPP is a beneficiary of the “National Council for Scientific and Technological Development” (CNPq) productivity fellowship. BPC received a PhD scholarship from the São Paulo Research Foundation (FAPESP; #2019/22511-3) during the development of this work and is currently supported by a FAPESP postdoctoral fellowship (FAPESP, #2025/08387-9).

### Competing interests

The authors have no conflict of interest to declare.

### Data availability

The data of the present work will be available at request to the corresponding author under reasonable conditions.

### Authors’ contributions statement

BPC: Conceptualization, data curation, formal analysis, funding acquisition, investigation, methodology, visualization, writing – original draft, writing – review & editing; VRB: Formal analysis, methodology, writing – original draft. AMR: Funding acquisition, resources, writing – review & editing. LPP: Funding acquisition, resources, writing – review & editing.; DE: Conceptualization, funding acquisition, methodology, resources, visualization, writing – review & editing.

### Ethics approval

The institutional ethics committee approved the present study, CEUA application n° 9512270320, certifiable at http://ceua.sirpp.unifesp.br/.

### Consent to participate

Not applicable.

### Consent for publication

Not applicable.

## Notes

### Competing Interest Statement

The authors have declared no competing interest.

### Summary of Updates

Revisions were carried out as part of the peer review process of Brain and Behavior (ISSN 2162-3279). The manuscript has now been approved and will be available at DOI:10.1002/brb3.71492

