## Supplementary material for "Rapid metabolic and behavioural maladaptations following short-term obesogenic diet withdrawal during post-weaning development in male Wistar rats": Suplementary table

**Supplementary table 1. Model information on single observation data parameters: body mass, adiposity, and metabolic.**

| Dependant variable | Model information |
|---|---|
| Initial body mass | $\chi^2 = 0.28$, p = 0.866, rDf = 12, $r^2 = 0.02$, pw = 0.068 |
| Final body mass | $\chi^2 = 0.13$, p = 0.936, rDf = 12, $r^2 = 0.01$, pw = 0.059 |
| Body mass gained (%) | $\chi^2 = 1.93$, p = 0.381, rDf = 12, $r^2 = 0.13$, pw = 0.202 |
| Adiposity | |
| Visceral fat (g/100g) | $\chi^2 = 61.94$, p < 0.001, rDf = 12, $r^2 = 0.84$, pw > 0.999 |
| Mesenteric fat (g/100g) | $\chi^2 = 20.04$, p < 0.001, rDf = 12, $r^2 = 0.63$, pw = 0.982 |
| Epididymal fat (g/100g) | $\chi^2 = 98.09$, p < 0.001, rDf = 12, $r^2 = 0.89$, pw > 0.999 |
| Retroperitoneal fat (g/100g) | $\chi^2 = 44.43$, p < 0.001, rDf = 12, $r^2 = 0.79$, pw > 0.999 |
| Serum | |
| Glucose (mg/dL) | $\chi^2 = 19.39$, p < 0.001, rDf = 12, $r^2 = 0.62$, pw =0.979 |
| Insulin (ng/mL) | $\chi^2 = 35.10$, p < 0.001, rDf = 12, $r^2 = 0.74$, pw > 0.999 |
| HOMA-IR index | $\chi^2 = 23.93$, p < 0.001, rDf = 12, $r^2 = 0.67$, pw =0.994 |
| Corticosterone (ng/mL) | $\chi^2 = 32.10$, p < 0.001, rDf = 1.5, $r^2 = 0.73$, pw = 0.999 |
| Triacylglycerol (mg/dL) | $\chi^2 = 338.94$, p < 0.001, rDf = 12, $r^2 = 0.97$, pw > 0.999 |
| Cholesterol (mg/dL) | $\chi^2 = 3.45$, p = 0.178, rDf = 12, $r^2 = 0.22$, pw = 0.350 |
| HDL cholesterol (mg/dL) | $\chi^2 = 16.75$, p < 0.001, rDf = 12, $r^2 = 0.58$, pw = 0.955 |
| n-HDL cholesterol (mg/dL) | $\chi^2 = 2.21$, p = 0.330, rDf = 12, $r^2 = 0.16$, pw = 0.248 |
| Liver | |
| Glycogen (mg/g) | $\chi^2 = 47.75$, p < 0.001, rDf = 12, $r^2 = 0.87$, pw > 0.999 |
| Triacylglycerol (mg/100g) | $\chi^2 = 18.02$, p < 0.001, rDf = 12, $r^2 = 0.61$, pw = 0.973 |
| Cholesterol (mg/100g) | $\chi^2 = 1.06$, p = 0.587, rDf = 12, $r^2 = 0.08$, pw = 0.135 |
| HDL cholesterol (mg/100g) | $\chi^2 = 2.11$, p = 0.348, rDf = 12, $r^2 = 0.15$, pw = 0.232 |
| n-HDL cholesterol (mg/100g) | $\chi^2 = 0.90$, p = 0.637, rDf = 12, $r^2 = 0.07$, pw = 0.123 |

(rDf) residual degrees of freedom; (pw) power. Samples size = 5/group

**Supplementary table 2. Model information on single observation data parameters: inflammation and oxidative stress.**

| Dependant variable | Model information |
|---|---|
| Serum | |
| IL6 | $\chi^2 = 0.17$, $p = 0.917$, rDf = 12, $r^2 = 0.01$, pw = 0.059 |
| IL10 | $\chi^2 = 0.21$, $p = 0.898$, rDf = 12, $r^2 = 0.01$, pw = 0.059 |
| TNFα | $\chi^2 = 13.56$, $p = 0.001$, rDf = 12, $r^2 = 0.53$, pw = 0.908 |
| IL1β | $\chi^2 = 31.20$, $p < 0.001$, rDf = 12, $r^2 = 0.70$, pw = 0.998 |
| Hypothalamus | |
| IL6 | $\chi^2 = 19.87$, $p < 0.001$, rDf = 12, $r^2 = 0.62$, pw = 0.979 |
| IL10 | $\chi^2 = 8.77$, $p = 0.012$, rDf = 12, $r^2 = 0.42$, pw = 0.739 |
| TNFα | $\chi^2 = 12.36$, $p = 0.002$, rDf = 12, $r^2 = 0.51$, pw = 0.883 |
| IL1β | $\chi^2 = 52.54$, $p < 0.001$, rDf = 12, $r^2 = 0.81$, pw > 0.999 |
| Liver | |
| IL6 | $\chi^2 = 14.04$, $p = 0.009$, rDf = 12, $r^2 = 0.54$, pw = 0.929 |
| IL10 | $\chi^2 = 24.13$, $p < 0.001$, rDf = 12, $r^2 = 0.67$, pw = 0. 995 |
| TNFα | $\chi^2 = 1.74$, $p = 0.420$, rDf = 12, $r^2 = 0.13$, pw = 0.200 |
| IL1β | $\chi^2 = 1.69$, $p = 0.430$, rDf = 12, $r^2 = 0.12$, pw = 0.196 |
| Hippocampus | |
| IL6 | $\chi^2 = 18.82$, $p < 0.001$, rDf = 12, $r^2 = 0.61$, pw = 0.973 |
| IL10 | $\chi^2 = 1.88$, $p = 0.390$, rDf = 12, $r^2 = 0.14$, pw = 0.217 |
| TNFα | $\chi^2 = 19.04$, $p < 0.001$, rDf = 12, $r^2 = 0.62$, pw = 0.979 |
| IL1β | $\chi^2 = 7.81$, $p = 0.020$, rDf = 12, $r^2 = 0.36$, pw = 0.623 |
| Mesenteric fat | |
| IL6 | $\chi^2 = 2.93$, $p = 0.231$, rDf = 12, $r^2 = 0.19$, pw = 0.297 |
| IL10 | $\chi^2 = 10.397$, $p = 0.004$, rDf = 12, $r^2 = 0.48$, pw = 0.841 |
| TNFα | $\chi^2 = 30.11$, $p < 0.001$, rDf = 12, $r^2 = 0.72$, pw = 0.999 |
| IL1β | $\chi^2 = 9.41$, $p = 0.009$, rDf = 12, $r^2 = 0.43$, pw = 0.757 |
| Hippocampus | |
| SOD | $\chi^2 = 48.46$, $p < 0.001$, rDf = 12, $r^2 = 0.81$, pw > 0.999 |
| CAT | $\chi^2 = 0.07$, $p = 0.954$, rDf = 12, $r^2 = 0.01$, pw = 0.059 |
| MDA | $\chi^2 = 15.33$, $p < 0.001$, rDf = 12, $r^2 = 0.58$, pw = 0.955 |
| CBP | $\chi^2 = 49.61$, $p < 0.001$, rDf = 12, $r^2 = 0.79$, pw > 0.999 |

(rDf) residual degrees of freedom; (pw) power. Samples size = 5/group

**Supplementary table 3. Cytokines absolute concentrations.**

| | | | |
|---|---|---|---|
| Serum | | | |
| IL6 | 105.921 ±20.317 | 97.767 ±32.846 | 99.498 ±39.961 |
| IL10 | 25.776 ±4.545 | 23.621 ±12.744 | 27.182 ±16.676 |
| TNFα | 21.536 ±1.012 | 22.541 ±0.724 | 23.665 ±0.969 |
| IL1β | 18.193 ±6.509 | 53.293 ±18.477 | 23.154 ±6.311 |
| Hypothalamus | | | |
| IL6 | 0.991 ±0.085 | 1.04 ±0.129 | 1.388 ±0.236 |
| IL10 | 0.725 ±0.074 | 0.887 ±0.108 | 0.76 ±0.088 |
| TNFα | 2.904 ±0.105 | 3.161 ±0.219 | 2.868 ±0.078 |
| IL1β | 0.316 ±0.031 | 0.141 ±0.034 | 0.285 ±0.051 |
| Liver | | | |
| IL6 | 5.058 ±1.049 | 5.927 ±1.301 | 3.586 ±0.446 |
| IL10 | 0.736 ±0.02 | 0.899 ±0.136 | 0.64 ±0.049 |
| TNFα | 2.328 ±0.235 | 2.402 ±0.083 | 2.279 ±0.061 |
| IL1β | 1.986 ±0.481 | 2.16 ±1.101 | 1.593 ±0.24 |
| Hippocampus | | | |
| IL6 | 1.087 ±0.096 | 1.522 ±0.285 | 1.484 ±0.154 |
| IL10 | 0.899 ±0.05 | 0.943 ±0.19 | 1.019 ±0.146 |
| TNFα | 2.822 ±0.443 | 3.119 ±0.214 | 3.724 ±0.167 |
| IL1β | 0.516 ±0.176 | 0.356 ±0.156 | 0.675 ±0.188 |
| Mesenteric fat | | | |
| IL6 | 4.624 ±1.95 | 3.328 ±0.585 | 4.103 ±1.086 |
| IL10 | 0.637 ±0.142 | 0.424 ±0.064 | 0.506 ±0.104 |
| TNFα | 2.441 ±0.123 | 2.306 ±0.103 | 2.751 ±0.165 |
| IL1β | 0.806 ±0.082 | 1.025 ±0.312 | 0.686 ±0.098 |

**Supplementary table 4. Model information on single observation data parameters: anxiety-like behaviour.**

| Dependant variable | Model information |
|---|---|
| Open field | |
| Lines crossed | $\chi^2 = 28.01$, $p < 0.001$, rDf = 12, $r^2 = 0.70$, pw = 0.999 |
| Time (Centre) | $\chi^2 = 12.32$, $p = 0.002$, rDf = 12, $r^2 = 0.52$, pw = 0.892 |
| Lines crossed (peripheral zone) | $\chi^2 = 30.69$, $p < 0.001$, rDf = 12, $r^2 = 0.69$, pw > 0.999 |
| Open Arms (EPM) | |
| Entries (% total entries) | $\chi^2 = 26.73$, $p < 0.001$, rDf = 12, $r^2 = 0.69$, pw = 0.997 |
| Time (% total time) | $\chi^2 = 13.82$, $p = 0.001$, rDf = 12, $r^2 = 0.54$, pw = 0.919 |
| Distance (% total distance) | $\chi^2 = 27.93$, $p < 0.001$, rDf = 12, $r^2 = 0.70$, pw = 0.998 |
| Average speed (m/s) | $\chi^2 = 0.05$, $p = 0.973$, rDf = 12, $r^2 = 0.00$, pw = 0.054 |
| Immobility (s) | $\chi^2 = 3.98$, $p = 0.137$, rDf = 12, $r^2 = 0.25$, pw = 0.414 |
| Enclosed Arms (EPM) | |
| Entries (% total entries) | $\chi^2 = 1.31$, $p = 0.519$, rDf = 12, $r^2 = 0.10$, pw = 0.160 |
| Time (% total time) | $\chi^2 = 4.02$, $p = 0.134$, rDf = 12, $r^2 = 0.25$, pw = 0.405 |
| Distance (% total distance) | $\chi^2 = 4.69$, $p = 0.096$, rDf = 12, $r^2 = 0.28$, pw = 0.463 |
| Average speed (m/s) | $\chi^2 = 8.67$, $p = 0.013$, rDf = 12, $r^2 = 0.42$, pw = 0.754 |
| Immobility (s) | $\chi^2 = 9.28$, $p = 0.010$, rDf = 12, $r^2 = 0.44$, pw = 0.784 |
| Centre (EPM) | |
| Entries (% total entries) | $\chi^2 = 12.35$, $p = 0.002$, rDf = 12, $r^2 = 0.51$, pw = 0.883 |
| Time (% total time) | $\chi^2 = 17.20$, $p < 0.001$, rDf = 12, $r^2 = 0.59$, pw = 0.962 |
| Distance (% total distance) | $\chi^2 = 22.08$, $p < 0.001$, rDf = 12, $r^2 = 0.65$, pw = 0.989 |
| Average speed (m/s) | $\chi^2 = 0.97$, $p = 0.617$, rDf = 12, $r^2 = 0.08$, pw = 0.130 |
| Immobility (s) | $\chi^2 = 0.64$, $p = 0.726$, rDf = 12, $r^2 = 0.05$, pw = 0.101 |
| Final third of the OA (EPM) | |
| Entries (% open arms) | $\chi^2 = 11.64$, $p = 0.003$, rDf = 12, $r^2 = 0.49$, pw = 0.855 |
| Time (% open arms) | $\chi^2 = 53.21$, $p < 0.001$, rDf = 12, $r^2 = 0.82$, pw > 0.999 |
| Distance (% open arms) | $\chi^2 = 72.87$, $p < 0.001$, rDf = 12, $r^2 = 0.86$, pw > 0.999 |
| Risk assessment | |
| Protected head dips (s) | $\chi^2 = 8.48$, $p = 0.014$, rDf = 12, $r^2 = 0.41$, pw = 0.744 |
| Unprotected head dips (s) | $\chi^2 = 16.02$, $p < 0.001$, rDf = 12, $r^2 = 0.57$, pw = 0.957 |
| Unprotected head dips (% total head dips | $\chi^2 = 11.21$, $p = 0.004$, rDf = 12, $r^2 = 0.48$, pw = 0.853 |
| Protected head dips (% protected time) | $\chi^2 = 14.58$, $p < 0.001$, rDf = 12, $r^2 = 0.53$, pw = 0.928 |
| Unprotected head dips (% unprotected time) | $\chi^2 = 1.16$, $p = 0.561$, rDf = 12, $r^2 = 0.08$, pw = 0.133 |
| Anxiety-like behaviour (PCA) | $\chi^2 = 20.07$, $p < 0.001$, rDf = 12, $r^2 = 0.63$, pw = 0.985 |

(rDf) residual degrees of freedom; (pw) power. Samples size = 5/group

**Supplementary table 5. 95% Confidence interval for correlations between inflammation, oxidative stress, and the open arms parameters.**

| | Serum Cort. | Serum TNFα | Hepatic IL6 | Ht IL6 | Hc TNF | M.FAT TNFα | Hc SOD | Hc MDA |
|---|---|---|---|---|---|---|---|---|
| Centre (OF) | | | | | | | | |
| Time (s) | | | 0.286 to 0.891 | | | −0.812 to −0.003 | | |
| Open Arms (EPM) | | | | | | | | |
| Entries (% total entries) | −0.931 to −0.489 | −0.861 to −0.163 | | −0.892 to −0.290 | −0.813 to −0.003 | −0.918 to −0.416 | −0.903 to −0.342 | −0.818 to −0.019 |
| Time (% total time) | −0.886 to −0.266 | | 0.336 to 0.902 | −0.894 to −0.299 | | | | |
| Distance (% total distance) | −0.882 to −0.249 | | 0.227 to 0.877 | −0.936 to −0.520 | | −0.841 to −0.092 | −0.913 to −0.391 | |
| Final third of the OA (EPM) | | | | | | | | |
| Time (% open arms) | −0.874 to −0.216 | −0.890 to −0.281 | | −0.896 to −0.312 | −0.899 to −0.325 | −0.836 to −0.078 | −0.899 to −0.325 | −0.834 to −0.069 |
| Distance (% open arms) | −0.917 to −0.411 | | 0.513 to 0.935 | −0.929 to −0.476 | −0.847 to −0.113 | −0.889 to −0.276 | −0.943 to −0.561 | −0.876 to −0.225 |
| Unprotected head dips (EPM) | −0.906 to −0.357 | −0.832 to −0.063 | 0.152 to 0.858 | −0.950 to −0.603 | | −0.879 to −0.234 | −0.863 to −0.173 | |
| ANX (PCA) | 0.257 to 0.884 | | −0.891 to −0.289 | 0.484 to 0.930 | | 0.866 to 0.184 | 0.350 to 0.905 | |

Cort: corticosterone; Hc: hippocampal; Ht: hypothalamic; M.FAT: mesenteric fat; SOD: superoxide dismutase; MDA: malondialdehyde * indicates $p < 0.05$
